# Impact of multi-echo ICA modeling decisions on motor-task fMRI analysis

**DOI:** 10.1101/2025.11.25.690440

**Authors:** Neha A. Reddy, Michelle C. Medina, Jordan N. Northrop, Chendong Zou, Rebecca G. Clements, Aravind Nehrujee, Milap Sandhu, Molly G. Bright

## Abstract

Multi-echo independent component analysis (ME-ICA) has been demonstrated to improve sensitivity and reliability of task functional magnetic resonance imaging (fMRI) data and, in particular, motor-task data with inherent task-correlated head motion. However, previous work has shown that an overly aggressive ME-ICA denoising approach may unintentionally remove task-related signal, while a more conservative approach may not effectively mitigate noise. While the effects of varied implementations of ME-ICA on signal and noise characteristics have been tested thoroughly in breath-hold data, the effects of similar modeling decisions have not been studied in motor-task data, which present with a more localized neural response. Here, we tested and compared the impacts of three analysis methods using rejected ME-ICA components as regressors in subject-level modeling: Aggressive (simple inclusion of ME-ICA regressors), Moderate (excluding task-correlated ME-ICA regressors from the model), and Conservative (orthogonalization of ME-ICA regressors to the core model and accepted ME-ICA components). We applied these methods to data from healthy and multiple sclerosis populations that included performance of hand-grasp, shoulder-abduction, and ankle-flexion tasks. We found that when the amount of head motion and its correlation with the task was high and the expected task-evoked signal was relatively low, the Conservative method led to significantly higher activation, t-statistics, and test-retest similarity in motor regions compared to the Aggressive method. Future motor-task studies may wish to implement similar models to prevent loss of motor signal, while still mitigating the effects of task-correlated head motion.

## Introduction

Multi-echo independent component analysis (ME-ICA) has demonstrated significant benefits in processing of blood oxygenation level dependent (BOLD) functional magnetic resonance imaging (fMRI) data during task performance. ME-ICA involves acquisition of fMRI data at multiple echo times, optimal combination of these echo times to improve signal and contrast, and use of ICA in denoising. Calculated kappa and rho parameters from the multiple echoes help to differentiate BOLD from non-BOLD ICA components (Kundu et al., 2012, 2013). Across various task-fMRI datasets, ME-ICA was found to improve sensitivity and reliability and decrease artifacts and inter-subject variability in task-fMRI studies, compared to the use of standard single-echo methods (Cohen et al., 2018; Cohen, Jagra, Visser, et al., 2021; Cohen, Jagra, Yang, et al., 2021; Cohen & Wang, 2019; Evans et al., 2015; Gonzalez-Castillo et al., 2016; Lombardo et al., 2016; Moia et al., 2021; Reddy et al., 2024).

In motor-task data, the common fMRI confound of head motion is additionally complicated by task-correlation of the head motion (Reddy et al., 2024). Our previous work demonstrated that ME-ICA mitigated artifacts and improved reliability of activation estimates in a hand-grasp dataset collected in healthy participants (Reddy et al., 2024). In this work, we incorporated rejected ME-ICA components (i.e., components classified as noise) as nuisance regressors in subject-level models. This aggressive denoising approach was effective at mitigating artifacts even in datasets with high levels of task-correlated head motion.

However, this implementation of ME-ICA may have limitations when applied to other types of motor-task data. Varied motor tasks have been associated with variable levels of activation given the same fMRI scan length, likely influenced by the number of muscle fibers involved in the movement (Rangaprakash et al., 2025). A hand-grasp task involves several dexterous muscles, and therefore a high number of muscle fibers and larger cortical representation, compared to a more isolated movement, such as ankle plantar flexion. As hand grasp should also lead to relatively strong motor activation with fMRI, an effective denoising method for hand-grasp motor data may not be directly applicable to all motor tasks. In our own group’s initial testing, we observed that an aggressive method of denoising limited our ability to detect activation in a shoulder movement dataset. In addition, introducing variable motor tasks and participants with clinical conditions may affect motion characteristics of the data and the impact of ME-ICA methods on subsequent denoising (Reddy et al., 2024; Seto et al., 2001). Hence, the specific implementation of ME-ICA components in fMRI modeling may need to consider the type of motor task and the population involved.

One approach to addressing the issue of signal loss with ME-ICA denoising was explored by Moia and colleagues (Moia et al., 2021), who tested the impacts of various implementations of rejected ME-ICA components in subject-level modeling of breath-hold task data with strongly task-correlated motion artifacts. In addition to an aggressive denoising approach similar to that implemented in our previous work (Reddy et al., 2024), the authors tested alternative denoising models to explore the trade-off between noise reduction and retention of the signal of interest. Moia and colleagues found that their conservative model led to the most reliable vascular activity estimates. This conservative model involved orthogonalization of rejected ME-ICA components to accepted ME-ICA components, the task regressor, motion regressors, and Legendre polynomials. However, this strategy has not been tested in motor-task data: whereas breath-holding evokes a systemic vasodilatory BOLD fMRI response, motor tasks evoke localized regions of neural activation and may also induce even greater head motion artifacts. While conservative orthogonalization may provide improvements to motor-task data analysis, the advantages and shortcomings of its utilization for this purpose are not known.

Orthogonalization is one method of dealing with unwanted collinearity in a model, which ideally should be minimized (Mumford et al., 2015). Motor tasks inherently cause motion that is correlated with the task and lead to task-correlated regressors, including task-correlated ME-ICA components. Therefore, another potential method to explore is the exclusion of task-correlated ME-ICA components from the regression model, which may be able to reduce collinearity and aid in retention of the desired motor signal. While this strategy prevents task-related signal from being attributed to highly collinear ME-ICA nuisance regressors, it is not as conservative as orthogonalization: orthogonalization ensures that *none* of the ME-ICA nuisance regressors share variance with the task model, other nuisance regressors, or any of the ME-ICA components demonstrating BOLD-weighting.

In this study, we sought to evaluate the impact of ME-ICA modeling decisions on data quality, motor activation estimates, and test-retest similarity in motor-task datasets. We compare three analysis methods using rejected ME-ICA components as nuisance regressors in subject-level modeling: Aggressive (simple inclusion of ME-ICA regressors in the design matrix), Moderate (exclusion of task-correlated ME-ICA regressors from the design matrix), and Conservative (orthogonalization of ME-ICA regressors to the core model and accepted ME-ICA components). We test each of these methods in four different datasets to understand the effects of these modeling techniques on data with variable motor tasks, motion characteristics, and participant task performance. These datasets include hand-grasp and shoulder-abduction tasks in healthy individuals and hand-grasp and ankle-flexion tasks in individuals with multiple sclerosis. Ultimately, these datasets allow us to comprehensively evaluate our three analysis methods and provide recommendations for robust mapping of motor activation in diverse datasets.

## Methods

### Participants and scan protocols

Four motor-task datasets were included in the analysis and are described in detail below. Functional MRI scan parameters for each dataset are summarized in Supplemental Table 1.

#### Healthy hand-grasp dataset (HealthyHand)

This study was approved by the Northwestern University Institutional Review Board, and all participants provided written, informed consent. Eight right-handed, healthy participants with no history of neurological or vascular disorders (4M, 26 ± 2 years) underwent scanning on a 3T Prisma MRI system with a 64-channel head coil. This dataset was previously described in Reddy et al. 2024 (Reddy et al., 2024).

Hand-grasp motor task: As described previously (Reddy et al., 2024), participants performed an isometric unimanual hand-grasp task at 40% maximum force. Each task trial was a 10-s "squeeze" and 15-s "relax", alternating 4 trials per hand, for a total of 16 trials per functional scan. Participants viewed task instructions and real-time force feedback on a screen, with a target box of 35-45% maximum force indicating the target force level. During the functional scans, force data from both hands were recorded with load cells (Interface, Inc.), at a sampling rate of 20 Hz.

Participants performed the hand-grasp motor task during two functional scans in the same scan session using a multi-band multi-echo gradient-echo echo planar imaging sequence provided by the Center for Magnetic Resonance Research (CMRR, Minnesota): TR = 2 s, TEs = 10.8/28.03/45.26/62.49/79.72 ms, FA = 70°, MB factor = 4, GRAPPA = 2, voxel size = 2.5x2.5x2 mm^3^, 210 volumes (Moeller et al., 2010; Setsompop et al., 2012). During the first functional scan (Limited), participants were instructed to keep their head as still as possible. During the second functional scan (Amplified), participants were instructed to add a small downwards nod at the beginning of each hand grasp and a small upwards nod to return to the starting position upon release of each hand grasp; this instruction added self-directed task-correlated motion to the Amplified motion scan.

#### Healthy shoulder-abduction dataset (HealthyShoulder)

This study was approved by the Northwestern University Institutional Review Board, and all participants provided written, informed consent. Six right-handed, healthy participants with no history of neurological or vascular disorders (3M, 23 ± 1 years) underwent scanning on a 3T Prisma MRI system with a 32-channel head coil.

Shoulder-abduction motor task: Participants performed an isometric unimanual shoulder-abduction task with their left arm at Low (14 ± 2% maximum), Medium (26 ± 6% maximum), and High (57 ± 10% maximum) torque. Each task trial was a 10-s “lift” and 15-s “relax”, with 7 consecutive trials at each torque level, for a total of 21 trials per functional scan. Participants viewed task instructions and real-time torque feedback on a screen, with a target box of goal ± 5% maximum torque indicating the target torque level. During the functional scans, force and torque data in six degrees of freedom were recorded from the load cell (JR3, Inc), at a sampling rate of 1000 Hz. A passive auditory stimulus was presented simultaneously during motor-task periods, as part of a separate study.

Participants performed the shoulder-abduction motor task during two functional scans in the same scan session using a multiband multi-echo gradient-echo echo planar imaging sequence provided by the Center for Magnetic Resonance Research (CMRR, Minnesota): TR = 2.2 s, TEs = 13.4/39.5/65.6 ms, FA = 90°, MB factor = 2, GRAPPA = 2, voxel size = 1.731 x 1.731 x 4 mm^3^, 44 slices, phase encoding direction A >> P, field of view 180 mm, matrix size 104 x 104, and 250 volumes (Moeller et al., 2010; Setsompop et al., 2012). Axial slices were aligned perpendicular to the base of the fourth ventricle. A reverse-phase encode P >> A scan was acquired before each functional scan for use in distortion correction.

#### Multiple sclerosis: hand-grasp (MSHand) and ankle-flexion (MSFoot) datasets

This study was approved by the Northwestern University Institutional Review Board, and all participants provided written, informed consent. Seven participants with multiple sclerosis (3M, 55 ± 15 years, Expanded Disability Status Scale 5.6 ± 1.3) underwent scanning on a 3T Prisma MRI system with a 64-channel head coil. One subject (MS-06) was scanned with a 32-channel head coil due to size restrictions with the 64-channel head coil.

Hand-grasp motor task: Participants performed an isometric unimanual hand-grasp task at 40% maximum force. Each task trial was a 10-s "squeeze" and a jittered "relax" period ranging from 12 to 18 s, alternating 7 trials per hand, for a total of 14 trials per functional scan. Participants viewed task instructions and real-time force feedback on a screen, with a target box of 35-45% maximum force indicating the target force level. During the functional scans, force data from both hands were recorded with load cells (Interface, Inc.), at a sampling rate of 100 Hz.

Ankle-flexion motor task: Participants performed a unilateral ankle task during which they alternated plantar flexion and dorsiflexion movements. Each task trial was 20-s "flex and extend", paced at 1-s per foot movement, and a jittered "relax" period ranging from 10 to 18 s, alternating 7 trials per foot, for a total of 14 trials per functional scan. Participants viewed task instructions and foot movement timing on a screen.

Participants participated in two scan sessions, a minimum of four weeks apart. These scans were obtained as part of a larger clinical trial consisting of four fMRI sessions, with an associated intervention (clinical trial ID: NCT06276634). The two scan sessions analyzed here were acquired prior to the intervention to avoid any impact of the intervention on the data. During each scan session, participants performed the ankle-flexion task during one functional scan and the hand-grasp task during a subsequent functional scan. Functional scans were collected using a multi-band multi-echo gradient-echo echo planar imaging sequence provided by the Center for Magnetic Resonance Research (CMRR, Minnesota): TR = 1.5 s, TEs = 10.6/27.83/45.06/62.29/79.52 ms, FA = 70°, MB factor = 4, GRAPPA = 2, voxel size = 2.5x2.5x2.5 mm^3^, 258 volumes for hand grasp and 324 volumes for ankle flexion (Moeller et al., 2010; Setsompop et al., 2012). A reverse-phase encode P >> A scan was acquired before each functional scan for use in distortion correction.

#### All datasets

All participants underwent a structural T1-weighted multi-echo MPRAGE scan using parameters adapted from Tisdall and colleagues (Tisdall et al., 2016): TR=2.17 s, TEs=1.69/3.55/5.41 ms, TI=1.16 s, FA=7°, FOV=256x256 mm^2^, and voxel size=1x1x1 mm^3^. The three echo MPRAGE images were combined using root-mean-square. During all functional scans, CO_2_ was measured via a nasal cannula and gas analyzer (PowerLab, ADInstruments), at a sampling rate matching that of the force / torque recordings.

### Data analysis

#### Creation of motor task and end-tidal CO_2_ regressors

Task timings were convolved with a canonical hemodynamic response function, rescaled from 0 to 1, then demeaned. The demeaned task trace was then downsampled to the resolution of the functional MRI images.

As done previously, end-tidal CO*_2_* was calculated for use as a nuisance regressor in subject-level modeling (Reddy et al., 2024). Performance of motor tasks may cause participants to modify breathing patterns, which affects arterial CO*_2_* levels and brain vasodilation, independently from neural activity. End-tidal CO*_2_* can be used to non-invasively estimate arterial CO*_2_* (McSwain et al., 2010) and account for the effects of respiration on the fMRI signal. End-tidal peaks were detected in the CO*_2_* data using an automatic peak-finder in MATLAB, then manually inspected. The end-tidal peaks were interpolated to form an end-tidal CO*_2_* trace that was convolved with a canonical hemodynamic response function, rescaled to the range of the unconvolved timeseries, then demeaned. The demeaned trace was then downsampled to the resolution of the functional MRI images.

#### Structural MRI pre-processing

T1-weighted images for each subject were processed with FSL’s (Jenkinson et al., 2012) *fsl_anat*, which performs bias field correction and brain extraction.

#### Functional MRI pre-processing

FSL (Jenkinson et al., 2012) and AFNI (Cox J.S., 1996) tools were used for fMRI pre-processing.

The first 10 volumes of each echo were removed to allow the signal to achieve steady-state magnetization. Distortion correction was performed using *topup* (FSL) for all datasets other than HealthyHand, which did not collect reverse-phase encode scans. For HealthyShoulder data, each scan was divided into three segments by torque level, which were analyzed independently. For all functional scans, head-motion realignment was estimated for the first echo with reference to the Single Band reference image taken at the start of the scan (*3dvolreg*, AFNI) and then applied to all echoes (*3dAllineate*, AFNI). All images were brain extracted (*bet*, FSL).

*Tedana* (version 24.0.2) (Ahmed et al., 2024; Dupre et al., 2021) was used to calculate a T2*-weighted combination of the multi-echo datasets, producing the optimally combined (ME-OC) fMRI dataset. The ME-OC timeseries was converted to signal percentage change for further analysis. No spatial smoothing was performed.

Several nuisance regressors were generated from the fMRI data, including motion parameters from volume registration, motion parameter derivatives, and the average CSF timeseries. The CSF timeseries was calculated by segmenting the CSF from the T1-weighted image (fsl_anat, FSL) using a threshold of 50%, and then transforming this CSF segmentation to functional space (epi_reg, FLIRT, FSL). The ME-OC timeseries was averaged across all CSF voxels.

#### ME-ICA component classification

ME-ICA components were automatically classified as signal (accepted) or noise (rejected) based on the *demo_taskexternalregressors_model* included in *tedana* v24.0.2 (Ahmed et al., 2024). These parameters build on the widely used *tedana minimal* component classification steps, with an additional criterion assessing nuisance regressors. The classification parameters are also explained in more detail and documented on the *tedana* website. Based on the *tedana demo_taskexternalregressors_model*, components were rejected if:

a. Rho > kappa
b. Greater number of voxels fit to S0 decay model compared to T2* decay model
c. Dice overlap is greater for S0 decay model compared to T2* decay model
d. Noise > signal
e. Kappa < kappa elbow
f. Rho ≥ rho elbow and kappa ≤ 2*rho
g. F test for nuisance regressors: p < 0.05 and R^2^ > 0.5

This approach generates large numbers of rejected components with low variance, and their inclusion in a denoising model would negatively impact degrees of freedom and potentially cause overfitting of the fMRI data. To mitigate this, components were ordered by their temporal variance. Components with variance < 0.1% were re-labeled as accepted (i.e., not rejected) until 1% of the total fMRI variance was accepted in this way.

#### Subject-level modeling

Each functional scan was modeled using a core model of 6 motion parameters and 6 motion derivatives from volume realignment, up to fourth-order Legendre polynomials, the end-tidal CO_2_ regressor, and motor-task regressor(s) (*3dREMLfit*, AFNI). Rejected ME-ICA components were incorporated into this core model using three different methods described below. Treatment of a representative ME-ICA component is demonstrated in Figure 1.

1. Aggressive: All rejected ME-ICA components were included in the subject-level model (e.g., added to the core model).
2. Moderate: As with (1), however, select rejected ME-ICA components were *not* included as regressors if they were correlated with the task regressor(s) (F test: P < 0.05 and R^2^ > 0.5 and kappa ≥ kappa elbow)
3. Conservative: All rejected ME-ICA components were orthogonalized to the accepted ME-ICA components, task regressor(s), motion regressors, Legendre polynomials, and end-tidal CO_2_ regressor, replicating the conservative model described by Moia and colleagues (2021), before inclusion in the subject-level model.

**Figure 1.**
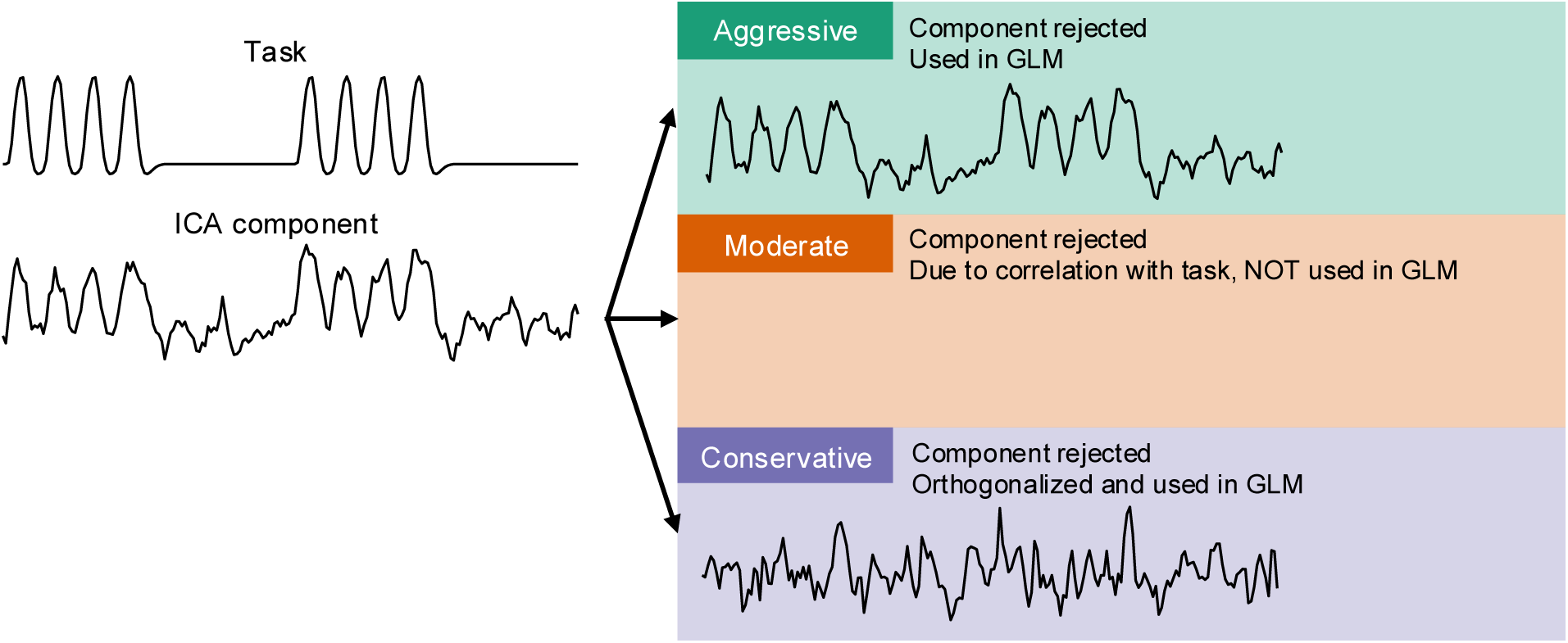
Treatment of a representative highly correlated ME-ICA component by Aggressive, Moderate, and Conservative models.

#### Quantification of head motion

Framewise displacement (FD) was calculated from the volume realignment parameters as the sum of the absolute difference of the realignment parameters between samples (Power et al., 2012). We investigated the correlation of the motion parameters with the task by calculating an adjusted R^2^ value for the model Task ∼ X + Y + Z + Roll + Pitch + Yaw. The task timing before convolution with canonical HRF was used here to match up the timing of motion and task performance. For tasks with right and left side movements, the two task timings were summed before implementation in the model to calculate motion correlation.

#### Motor ROI creation

Precentral gyrus and cerebellar ROIs were created for calculation of metrics within expected regions of motor activation. A precentral gyrus mask was created for each scan using the precentral gyrus mask, unthresholded, from the Harvard-Oxford Cortical Atlas (Desikan et al., 2006). The atlas region was transformed to each scan’s functional space using a concatenated transform of standard to anatomical space (*FLIRT* and *FNIRT*, FSL) and anatomical to functional space (*epi_reg*, *FLIRT*, FSL). A cerebellum mask was created for each scan using the MNI structural atlas region (Collins et al., 1995; Mazziotta et al., 2001) thresholded at 50%, then similarly transformed to each scan’s functional space.

For calculation of tSNR and spatial correlation, masks of the gray matter within the precentral gyrus and cerebellum were used. Each subject’s gray matter mask was calculated by segmenting the gray matter from the T1-weighted image (*fsl_anat*, FSL) and thresholding at 50%, then transforming this gray matter segmentation to functional space (*epi_reg*, *FLIRT*, FSL). The resultant ROI masks were created by finding the intersection of each subject’s gray matter mask with the precentral gyrus and cerebellum masks in functional space. For calculation of spatial correlation, the gray matter within the contralateral precentral gyrus and ipsilateral cerebellum were used as ROIs, as these hemispheres are most strongly associated with motor activity in the respective brain regions (Ashida et al., 2019; Penfield & Rasmussen, 1950).

For calculation of activation-related metrics (t-statistic, percent activated voxels), group-level analysis was used to identify the specific motor regions related to each task (e.g. hand, foot, shoulder). For each motor task and model (Aggressive, Moderate, Conservative), the beta coefficient and t-statistic maps from subject-level analysis were converted to MNI space by applying a concatenated transform of functional to anatomical space (*FLIRT* and *FNIRT*, FSL) and anatomical to functional space (*epi_reg*, *FLIRT*, FSL). Group-level analysis was performed with AFNI’s *3dMEMA* (Chen et al., 2012), using contrasts for each motor task > 0. Group-level maps were thresholded at p < 0.005 and clustered at alpha < 0.05, using a right-sided t-test (*3dFWHMx*, *3dClustSim*, *3dClusterize*, AFNI). The union of the resultant clusters from the Aggressive, Moderate, and Conservative models was considered to be the group-level region of significant motor activity. This union mask was transformed to each subject’s functional space. For each motor task, the union mask within the contralateral upper-extremity (hand and shoulder tasks) or lower-extremity (foot task) primary motor cortex (Fan et al., 2016) and ipsilateral cerebellum were used as ROIs.

#### Calculation of temporal signal-to-noise ratio (tSNR)

The temporal signal-to-noise ratio (tSNR) of the denoised models was compared to understand their effects on data quality. The residual timeseries from each model was used as the denoised dataset, and these residual timeseries were reverted from signal percentage change for each voxel to represent the true mean in tSNR analysis. Each voxel Y was converted as [Y*mean(X)]+mean(X), with X being the original voxel timeseries before signal percentage change conversion. tSNR of the denoised datasets was calculated for each voxel by dividing the mean of the timeseries by the standard deviation. A median tSNR was found within masks of the precentral gyrus and cerebellum grey matter. Significant differences in the tSNR between the Aggressive, Moderate, and Conservative models were tested as tSNR ∼ Model + (1|Subject).

#### Calculation of activation metrics

Summary activation metrics for each scan were calculated by analyzing the t-statistic and percent activated voxels within the motor ROIs; here, the primary motor cortex union mask created from group-level analysis was used as the ROI. The median t-statistic for each scan was found within the motor ROIs. The number of positive activated voxels (voxels showing a statistical significance of p_FDR_ < 0.05) within the motor ROI was divided by the total number of voxels within the ROI to calculate the percent activated voxels for each scan. Significant differences in the activation metrics between the Aggressive, Moderate, and Conservative models were tested as Metric ∼ Model + (1|Subject).

#### Calculation of spatial correlation between repeated runs

Subjects performed two runs of each motor task, and the spatial correlation between the resulting beta coefficient maps was calculated to assess test-retest similarity. Voxel-wise spatial correlation analysis was performed within the precentral gyrus and cerebellum gray matter masks using AFNI’s *3ddot,* which calculated a correlation coefficient for each pair of repeated runs within each mask. Significant differences in the spatial correlation between the Aggressive, Moderate, and Conservative models were tested as Correlation ∼ Model + (1|Subject).

#### Division of scans into groups

In this study, we analyzed motor-task data from three motor tasks (hand grasp, ankle flexion, and shoulder abduction) and two participant groups (healthy, MS). To better understand the effects of the tested models on these variable datasets, we divided them into groups based on expected activation intensity and observed motion characteristics. In terms of activation intensity, hand-grasp data was considered to have “high signal” compared to ankle flexion and shoulder abduction (“low signal”) (Rangaprakash et al., 2025). We analyzed the FD and adjusted R^2^ values to understand the spread of motion characteristics across scans. The median FD and R^2^ values were found and used to separate scans into “high FD” and “low FD” groups and “high R^2^” and “low R^2^” groups. Three scans were found to have extremely high FD, and were separated into a third “extreme FD” group.

## Results

All structural and functional scans were successfully collected as described above. The HealthyShoulder Low torque task was excluded from subject-level analysis due to low activation in each individual scan, making scan-level activation metrics unreliable. During the ME-ICA component classification process, across all datasets, ME-ICA components were accepted due to low variance in four functional scans: MSHand MS-01 run2 (1 component), MSHand MS-05 run1 (20 components) and run2 (19 components), and MSFoot MS-01 run2 (7 components). In addition, in over half the datasets, no task-correlated ME-ICA components were identified; therefore, the Aggressive and Moderate models are equivalent in many cases. A summary of task-correlated components identified across datasets, and temporal and spatial information for two example task-correlated components are shown in Supplemental Figure 1. The task-correlated components showed high temporal similarity to the task regressors, and spatial maps demonstrated positive amplitude in regions of expected motor activity.

### Motion characteristics of datasets

The average FD and the adjusted R^2^ for each scan are shown in Figure 2A. Note that the motion characteristics between the repeated runs of HealthyHand are expected to be very different, as participants were instructed to increase their head motion during Run 2. The distributions of FD and R^2^ values across all scans are shown in Figure 2B. Both FD and R^2^ demonstrate a distribution with one major peak, with three scans having extreme FD values outside this main distribution. The median FD (0.29) and R^2^ (0.35) values that were used for creation of low and high motion groups are shown as vertical lines in Figure 2B.

**Figure 2.**
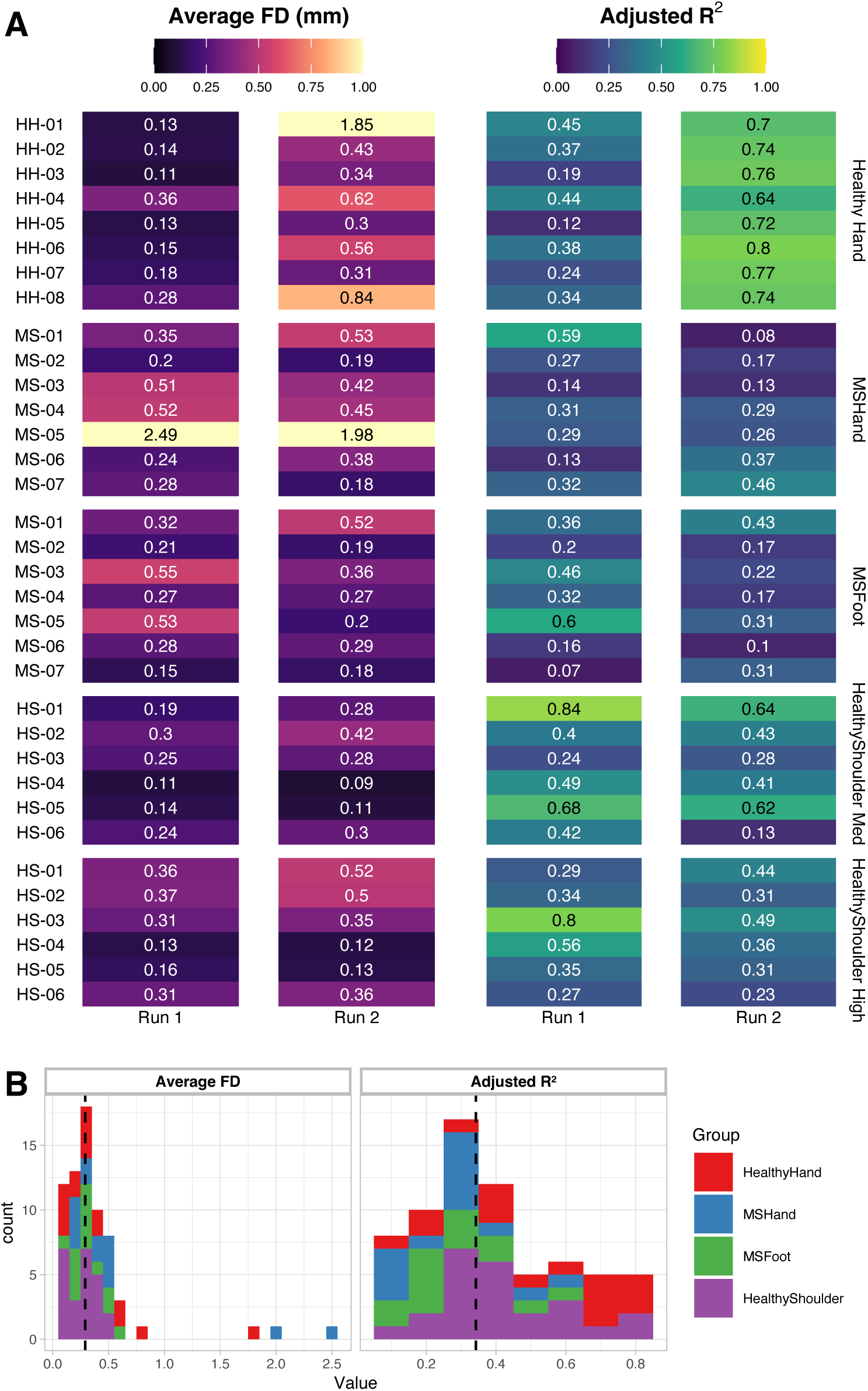
(A) Motion characteristics of all datasets: average Framewise Displacement (FD) across the scan and correlation of the motion parameters with the task (adjusted R^2^ value for the model Task ∼ X + Y + Z + Roll + Pitch + Yaw). (B) Distribution of FD and adjusted R^2^ values across all datasets. Dashed vertical line indicates median FD and adjusted R^2^ value.

### Temporal signal-to-noise ratio (tSNR)

tSNR was calculated for every scan to understand the impact of modeling decisions on overall data quality by summarizing residual unexplained variance from each modeling strategy. Median tSNR was calculated in gray matter of the precentral gyrus and cerebellum, where motor activity was expected (Figure 3). Across all datasets, the Conservative model had lower median tSNR than the Aggressive and Moderate models, with this difference reaching significance in the MSHand, MSFoot, and HealthyShoulder datasets. In the Conservative model, orthogonalization of the rejected ME-ICA components to the core model and all accepted (BOLD-weighted) ME-ICA components leads to reduced removal of fMRI timeseries variance, and therefore greater overall noise. The HealthyShoulder dataset had higher overall tSNR compared to the

**Figure 3.**
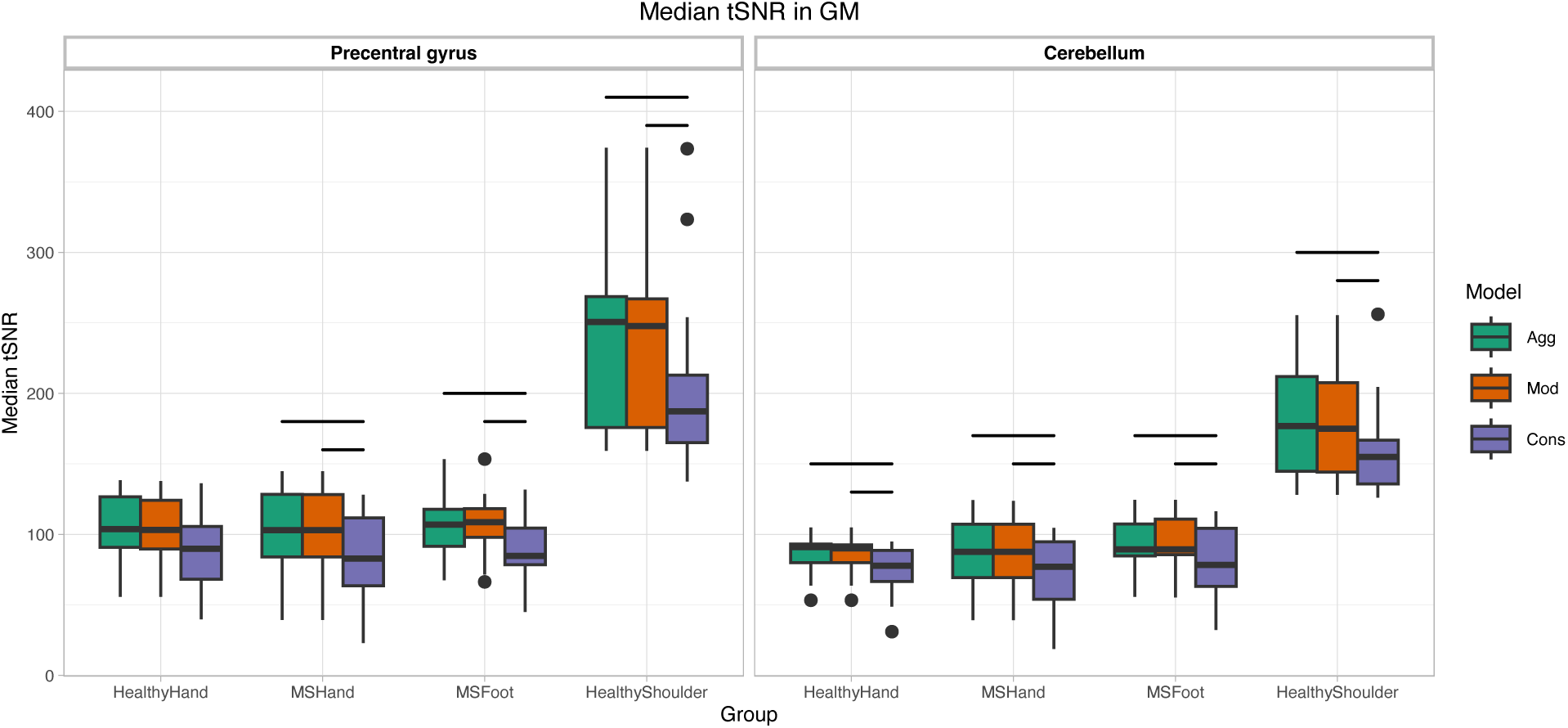
Median temporal signal-to-noise ratio (tSNR) for each dataset and model within gray matter (GM) of the precentral gyrus and cerebellum. Horizontal bars indicate significant differences between models (p < 0.05, Bonferroni-corrected).

### Group-level motor activation

We compared group-level activation maps to visualize broad patterns of activation in each dataset and understand the overall impacts of the Conservative, Moderate, and Aggressive models on these patterns (Figure 4). The HealthyHand, MSHand, and MSFoot datasets demonstrated strong motor activation in the precentral gyrus across all models. However, for the HealthyShoulder dataset, the Aggressive model reduced detection of precentral gyrus activation compared to the Moderate and Conservative models, which shows the potential importance of less aggressive denoising in group-level analysis.

**Figure 4.**
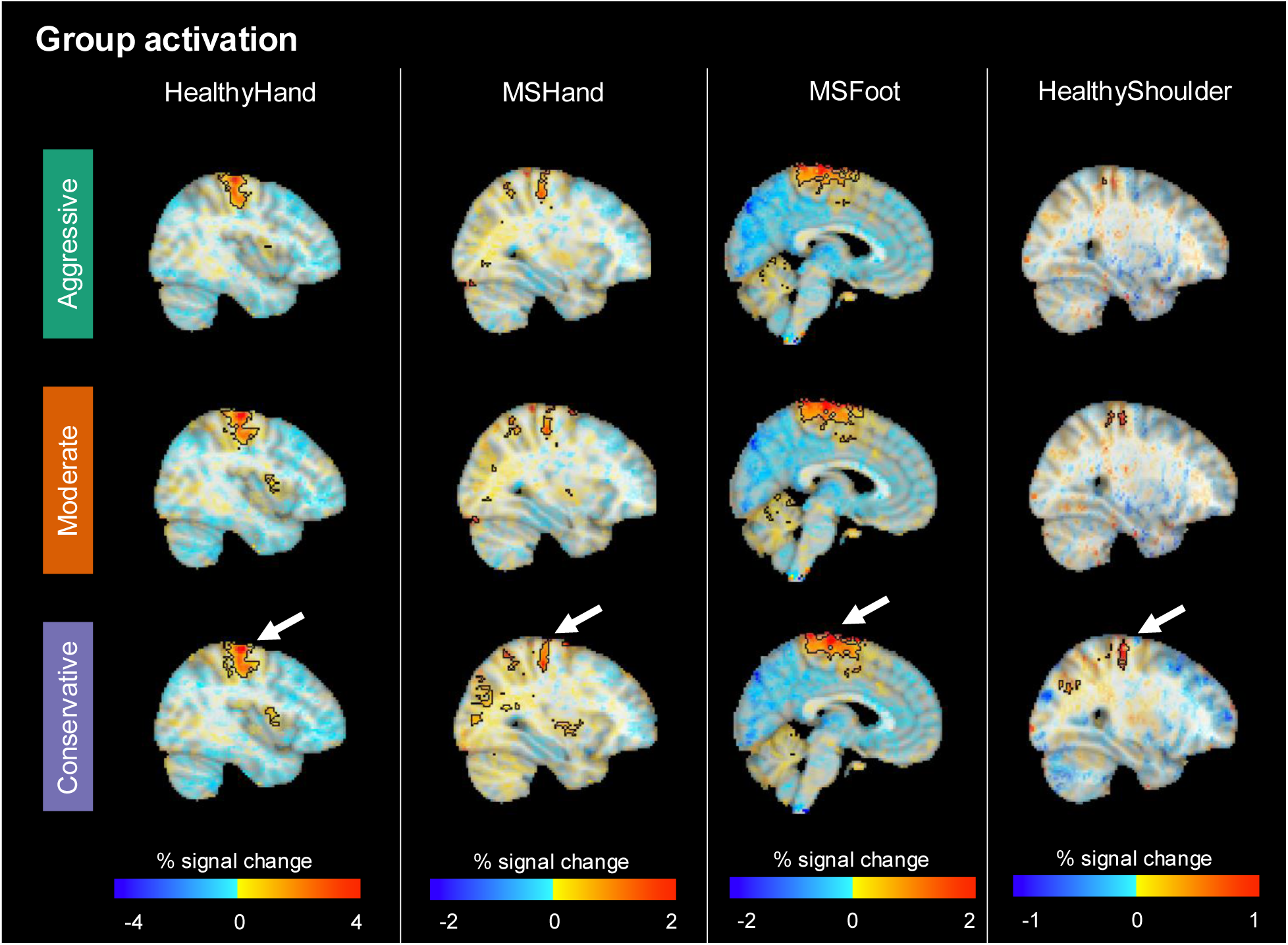
Group-level activation during a left-sided motor task for each dataset. Sagittal views of the right hemisphere are shown. Opacity of beta coefficients is modulated by the t-statistic. Significant activation (p < 0.005, alpha < 0.05) is shown in black outline. White arrows indicate regions of precentral gyrus activation.

### Subject-level activation metrics

We compared the t-statistics and percent activated voxels in the contralateral primary motor cortex and ipsilateral cerebellum across models. In order to focus on the specific motor regions involved in each task, we used group-level analysis across all scans in each dataset to determine the significant clusters of activation in the contralateral primary motor cortex and ipsilateral cerebellum to use as ROIs for each motor task. In addition, to better identify trends across our diverse scans with variable signal and noise properties, we separated scans into groups based on expected signal activation intensity and motion characteristics. Hand-grasp motor tasks were considered to have high signal, and foot and shoulder motor tasks were considered to have relatively low signal. The overall amount of motion (FD) and task-correlation of motion (adjusted R^2^) were used to further separate scans into low and high categories. Three datasets with FD > 1 mm were placed in an extreme FD group.

Across all groups, the Aggressive model led to the lowest t-statistics in the primary motor cortex (Figure 5A). In the high signal group, the Conservative model significantly increased t-statistics compared to Aggressive when the FD or adjusted R^2^ were high. In some instances, both Moderate and Conservative models led to similar improvements in t-statistics compared to the Aggressive model, as shown in the MSFoot example in Figure 6A. The extreme FD group had very low t-statistics across all models. When motion was lower, both Moderate and Conservative models demonstrated improvements compared to Aggressive, with a representative low motion HealthyHand scan shown in Figure 6A. In the low signal group, the Conservative and Moderate models significantly increased t-statistics compared to the Aggressive model when FD was high, with the Conservative model improving t-statistic compared to both models when adjusted R^2^ was high. However, for a few scans, use of the Conservative model led to a more negative median t-statistic (indicated with blue arrows in Figure 5A); an example MSFoot scan demonstrating this effect is shown in Supplemental Figure 2. The example scan shows that the Conservative model led to retention of a high-amplitude negative t-statistic artifact in non-motor regions across the brain, which influenced the median t-statistic calculation.

**Figure 5.**
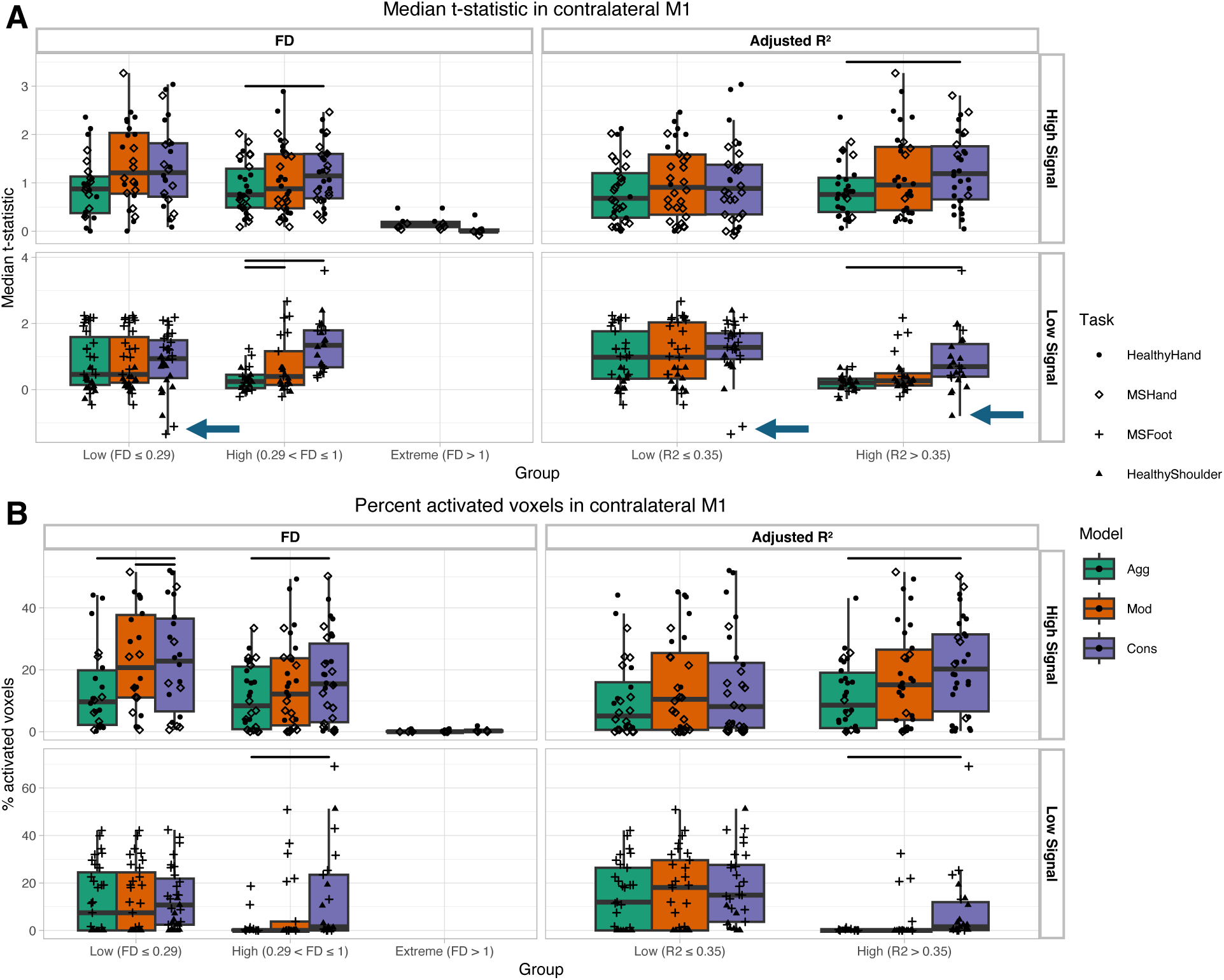
Activation metrics calculated in significant group-level regions within the contralateral primary motor cortex. Regions were determined as the union of significant regions across all three models. (A) Median t-statistic and (B) Percent activated voxels were calculated for each scan. Horizontal bars indicate significant differences between models (p < 0.05, Bonferroni-corrected). Blue arrows indicate scans in which use of the Conservative model led to a more negative t-statistic.

**Figure 6.**
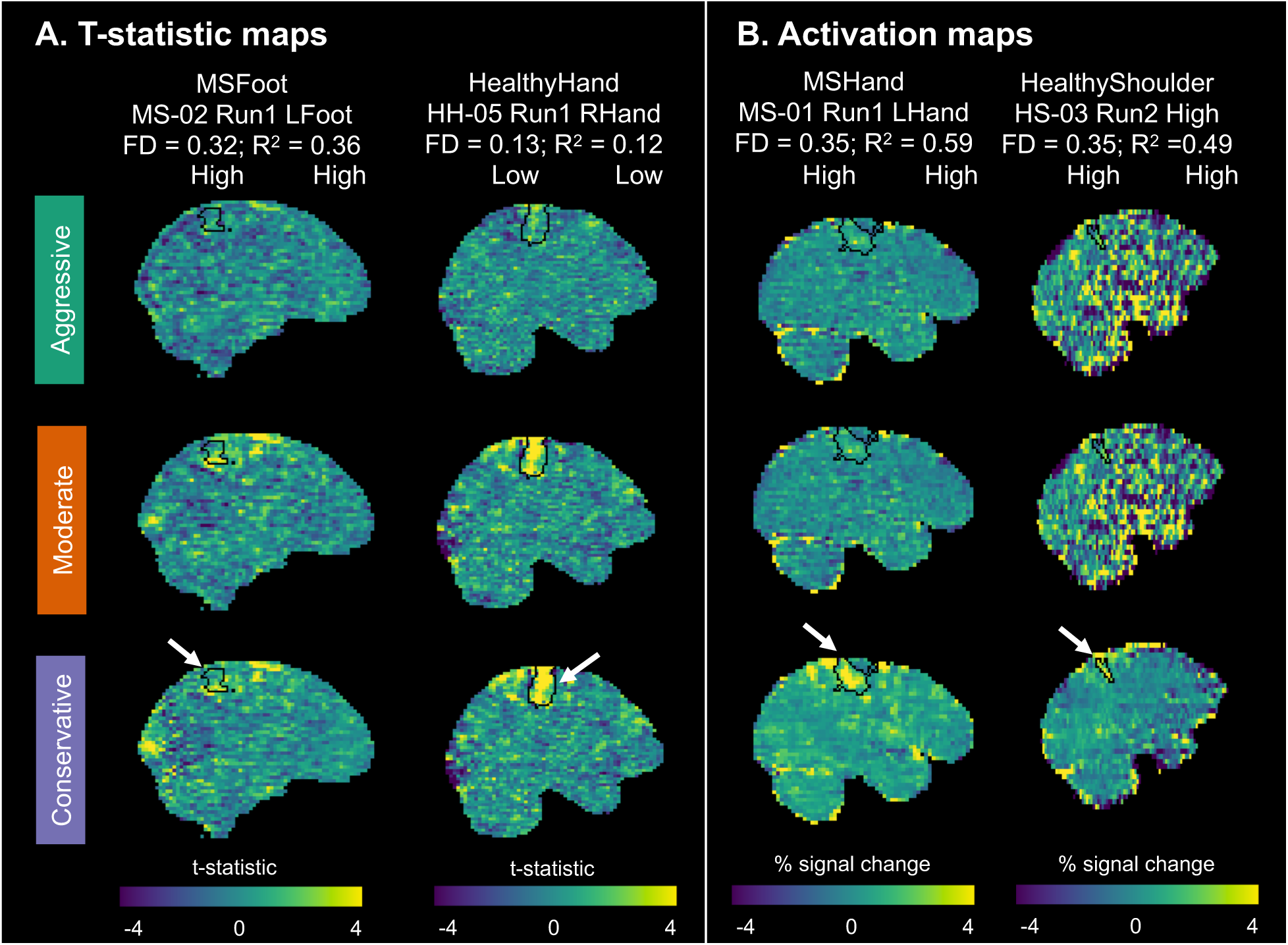
Example t-statistic and activation maps. The primary motor cortex ROI for each task is shown in black outline. (A) T-statistic maps: MSFoot and MSHand scans demonstrate similar t-statistics in motor region for Moderate and Conservative models. (B) Activation maps: MSHand and HealthyShoulder scans demonstrate expected motor activation with only the Conservative model.

The percent activated voxels were assessed to understand the impact of each model on the motor activation in the expected regions (Figure 5B). In the primary motor cortex, the Aggressive model led to the least activation compared to the Moderate and Conservative models. In the high signal group, the Conservative model significantly increased activation extent compared to Aggressive and Moderate models when FD was low and compared to the Moderate model when FD or adjusted R^2^ were high. In the low signal group, the Conservative model significantly increased activation extent compared to the Aggressive model when FD or adjusted R^2^ were high. Notably, in the high motion groups, the Aggressive and Moderate models led to zero or minimal activation in several low signal datasets, while the Conservative model allowed for visualization of motor activity. Representative high motion scans from the MSHand and HealthyShoulder groups demonstrate the increase in motor activation with the Conservative model (Figure 6B).

Similar t-statistic and activation trends were seen in the ipsilateral cerebellum. In this region, the Conservative model demonstrated significant improvements compared to the Aggressive model in several additional groups (Supplemental Figure 3). T-statistics for high signal datasets were significantly greater during low and high FD conditions and for low signal datasets when adjusted R^2^ was low. Percent activated voxels were also significantly improved with the Conservative model in the high signal group when FD was high.

### Spatial correlation between runs

We investigated spatial correlation between runs within each subject to understand the models’ impact on test-retest similarity of results. The average FD and adjusted R^2^ between the two runs were used to group scans. In the precentral gyrus gray matter, the high signal group demonstrated variable results across motion categories. The Conservative model outperformed the Aggressive and Moderate models in the high FD and low adjusted R^2^ groups, though not a significant trend. While this difference was significant in the extreme FD group, note that only activation maps from one subject were placed into this category after motion was averaged between runs. The Moderate model had higher spatial correlation that the Aggressive and Conservative models in the low FD and high adjusted R^2^ group.

In the low signal group, the Conservative model significantly improved spatial correlation compared to the Aggressive and Moderate models in every motion category. Figure 8 shows a representative HealthyShoulder scan with improved spatial correlation when implementing the Conservative model. However, one consequence of a Conservative denoising approach is retention of noise, shown by an extreme motion scan from the MSHand group (Figure 8). Though spatial correlation in precentral gyrus gray matter improved with the Conservative model, additional noise also remained throughout the brain. The Conservative model led to higher striping artifacts and apparent negative activation in extra-motor regions.

Similar trends were observed in the cerebellum, with some differences in statistical significance (Supplemental Figure 4). Spatial correlation in the cerebellum was significantly greater with the Conservative model compared to the other models when signal was high and FD was low. However, in the low signal group, no model demonstrated significant improvement over the others. This observation is largely due to a wider spread in spatial correlations using the Conservative model in the cerebellum

## Discussion

We tested the impact of three ME-ICA models on 56 fMRI scan runs from 21 participants, spanning three types of motor tasks and both healthy and MS populations. Hand motor tasks (“high signal” datasets) were expected to be associated with stronger neural activation compared to foot and shoulder tasks (“low signal” datasets). We found that the overall head motion and task-correlation of head motion varied widely across these scans, enabling a division of all scans into high and low motion groups. While data quality was lowest in the Conservative scans due to less aggressive denoising, the Conservative model led to higher t-statistics and greater activation in the motor regions of interest. This improvement was particularly pronounced in low signal datasets with high amounts of motion. Spatial correlation of activation patterns between repeated runs was also significantly higher using the Conservative model when signal was low.

### Motion characteristics of varied motor-task datasets

As expected, across our datasets, we observed variable degrees of overall motion and task-correlation of motion (Figure 2). Though ideally, experimental designs should be crafted to minimize head motion, our findings show that may not always be possible, particularly in studies involving larger amplitude movements and clinical populations. The FD thresholds used in this study (Low ≤ 0.29 mm, 0.29 mm < High ≤ 1 mm, Extreme > 1 mm) align with thresholds implemented in other studies. These studies have commonly used FD thresholds between 0.2 to 0.5 mm to censor high-motion data points (Afyouni & Nichols, 2018; Power et al., 2014). Our study’s threshold for High motion datasets falls within this range. Previous work has also demonstrated that post-stroke participants can exhibit head motion above 1 mm during performance of motor tasks (Reddy et al., 2024; Seto et al., 2001), aligning with our threshold for Extreme motion.

### Impact of modeling on activation metrics, test-retest similarity, and noise

In this study, we compared three methods of including ME-ICA nuisance regressors in subject-level models. The Aggressive model involved a simple inclusion of all rejected ME-ICA regressors as nuisance regressors in each subject-level model, as done previously in our work with the HealthyHand dataset (Reddy et al., 2024). The Moderate model sought to minimize model collinearity arising from task-correlated components and excluded any rejected ME-ICA components as regressors that were correlated with the motor task. The Conservative model aimed to prevent signal loss through orthogonalization of all rejected ME-ICA regressors to the core model and accepted ME-ICA components before inclusion in the subject-level model, as previously performed by Moia and colleagues for breath-hold data (Moia et al., 2021). We note that in many cases, the Moderate model was identical to the Aggressive model, as no rejected ME-ICA components were found to be correlated with the motor task.

We analyzed t-statistics and activation within the motor ROIs determined from group-level analysis to understand each model’s impact on motor activation findings. In high and low-signal data, the Conservative model outperformed the Aggressive model when overall motion or task-correlation of motion was high, leading to higher t-statistics and greater activation within the ROI (Figure 5, Supplemental Figure 3). The Conservative model also led to greater numbers of activated voxels in anticipated brain regions than the Moderate and Aggressive models in high-signal data when FD was low. In our few datasets with extreme motion, all three models recovered little motor activity.

As each of our subjects performed two runs of the same motor task, we calculated spatial correlation of the activation patterns between runs to understand the impact of the models on test-retest similarity (Figure 7, Supplemental Figure 4). The high-signal data demonstrated mixed results, with the Conservative model only significantly improving spatial correlation of extreme motion datasets. The Moderate model had significantly greater spatial correlation compared to the Aggressive model when task-correlation of motion was high. However, in the low-signal group, the Conservative model demonstrated significant improvements in spatial correlation compared to the other two models in every motion category. Previous work has shown that regression of motion and other denoising techniques can decrease reliability of results, while not necessarily reducing validity (Noble et al., 2019). Therefore, the residual effects of motion present with the Aggressive model may be contributing to greater spatial correlation between runs. While the spatial correlation results alone should not be used to decide between implementation of models as they do not directly reflect validity of results, they provide an additional perspective when comparing model outcomes.

**Figure 7.**
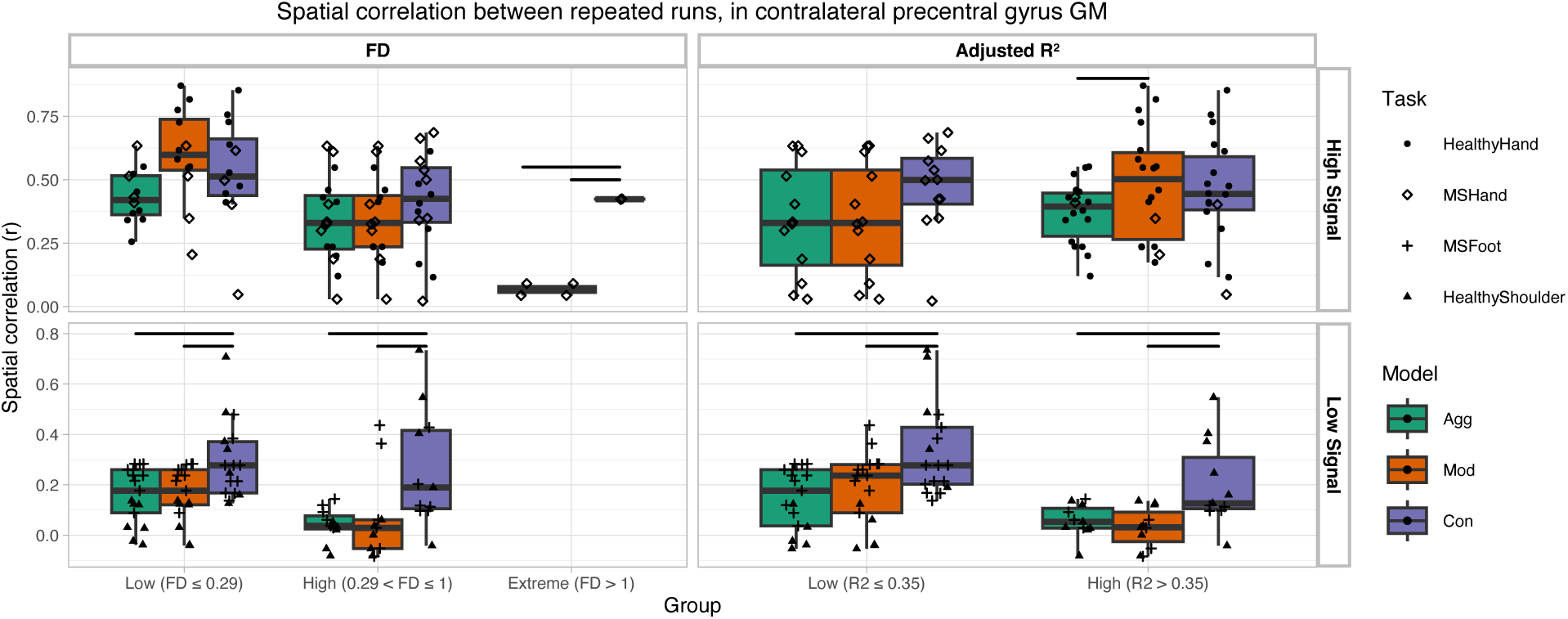
Spatial correlation between runs 1 and 2 for each dataset and model, calculated in gray matter (GM) of the contralateral precentral gyrus. Horizontal bars indicate significant differences between models (p < 0.05, Bonferroni-corrected).

Since the Conservative model prioritized the retention of signal related to the motor task, it also led to the highest remaining noise in the data. tSNR was significantly lower across datasets when using the Conservative model, compared to the Aggressive and Moderate models (Figure 3). In addition, activation maps from some individual subject datasets demonstrated greater noise and artifacts throughout the brain when using the Conservative model compared to the Aggressive and Moderate models (Figure 8). Therefore, analysis of non-motor regions with concurrent, more subtle activation may be affected by increased global noise when using the Conservative model.

**Figure 8.**
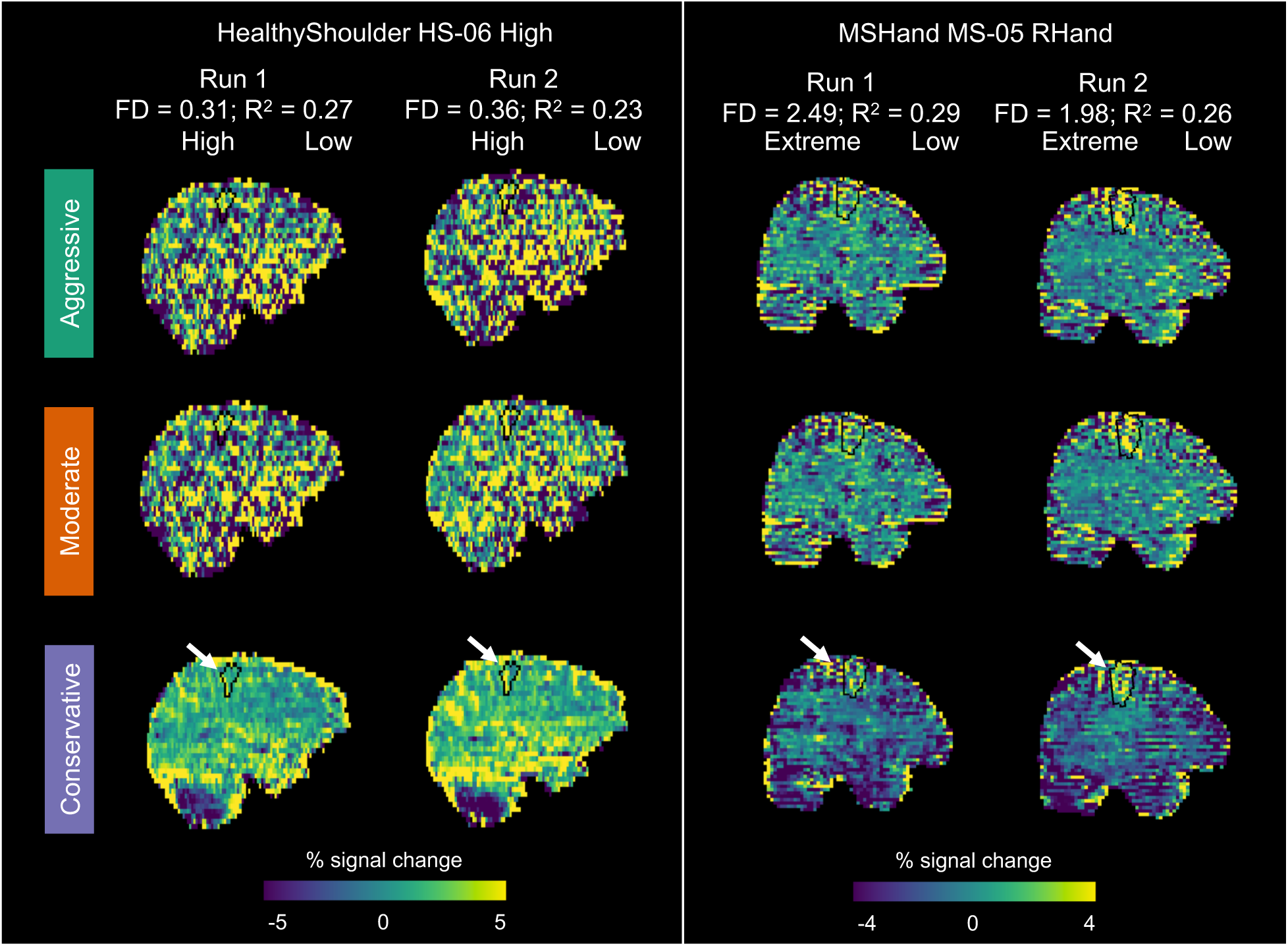
Example activation maps compared between runs. The primary motor cortex ROI for each task is shown in black outline to highlight the main region of activation. The HealthyShoulder scans demonstrate noise reduction and clearer identification of motor activity with the Conservative model. The MSHand scans show an example usage of the Conservative model that leads to retention of greater whole-brain noise when motion is extreme.

We note that while tSNR is often used as a measure of data quality (Murphy et al., 2007; Vizioli et al., 2021), the current results highlight the imperfect nature of this metric when comparing denoising techniques. Though the Aggressive and Moderate models led to higher tSNR due to greater overall variance explained by the models, this denoising also obscured motor-task signal, as some task-related variance was not attributed to the task regressors. The t-statistic results better reflect the balance between the detected task-related signal, residual model variance, and design matrix covariance. This nuance in interpreting tSNR may be important to consider in future comparisons of denoising strategies.

Group-level analysis showed that the Conservative model may be particularly beneficial in datasets with lower-signal motor tasks, like shoulder abduction (Figure 4). However, group-level results were similar across models in our HealthyHand, MSHand, and MSFoot datasets, demonstrating that an Aggressive denoising approach can be a useful group-level analysis tool in many datasets with strong noise artifacts and strong expected motor activation. This finding aligns with our previous hand-grasp results, which demonstrated high efficacy of an Aggressive ME-ICA denoising model in high-motion data (Reddy et al., 2024). However, motor activation in the HealthyShoulder dataset was strongly affected by Aggressive denoising, even at the group level, indicating that certain motor tasks may require a Conservative denoising approach for both subject- and group-level analyses.

### Recommendations for use of ME-ICA in motor-task studies

Our findings demonstrate that the Conservative model approach, involving orthogonalization of ME-ICA nuisance regressors to the core model and accepted ME-ICA components, significantly improves signal retention and activation metrics in motor-task fMRI data, particularly when motion is high and task-related signal is low. As it may be difficult to determine which motor tasks are high-signal without specific comparisons (Rangaprakash et al., 2025), the Conservative model can be an effective initial approach to analyzing motor-task data that “does no harm” in unintentionally removing signal. When analyzing a high-motion dataset, noise and artifact removal may be a particular concern. In these instances, the Conservative model can still be an appropriate approach if analysis is limited to motor regions in subject-level results or involves only group-level analysis. However, the Conservative model may also lead to retention of motion-related artifact throughout the brain. If subject-level analysis of extra-motor regions is desired and motion levels are high, the Moderate model may be preferred over the Conservative model to better mitigate noise, though the paucity of strongly task-correlated rejected ME-ICA components limits potential benefits of the Moderate model. Note that both the Aggressive and Moderate models may limit the ability to observe activation in primary motor regions in low-signal motor-task data. In extreme motion cases, even the Conservative model may not sufficiently identify the signal of interest (Figure 5). If extreme motion is present, experimenters may wish to implement additional noise mitigation during acquisition, use alternative denoising strategies, or modify the task protocol if motion-related degradation is too severe to be addressed by analysis techniques.

As future motor-task fMRI studies seek to understand motor activity in clinical populations with high head motion and lower signal due to potential weakness or impairment, usage of the Conservative model becomes particularly important to enable visualization of motor signal. Studies of proximal motor activity can also benefit from the Conservative model, especially as interest in motor-task fMRI broadens beyond hand grasp tasks (Gordon et al., 2023).

### Limitations and considerations

Though we included data from different participant populations and motor tasks, several other types of participants and tasks may affect motor activity and motion characteristics. To increase applicability of these results to a wider group of datasets, we categorized our scans by motor signal intensity and amount / task-correlation of motion. These categories may help inform future studies that understand the signal and motion characteristics of their own datasets.

In this study, we used preset task timings to create the task stimuli used in subject-level modeling. However, load cell recordings or other real-time assessments may improve modeling by accounting for precise magnitude and timing of motor activity and potentially help with collinearity issues. Real-time measurements may also affect identification of task-correlated ME-ICA components that were accepted in the Moderate model. Since the MSFoot task did not have real-time motor recordings available, we used preset task timing instructions across all motor tasks in this study to fairly compare results between tasks. As all models (Conservative, Moderate, and Aggressive) utilized the same task stimuli, this modeling decision should not substantially affect our model comparisons.

Additionally, we encourage future denoising studies to report both amount and task-correlation of head motion in their datasets, as we show that both metrics may be important for understanding the best denoising strategies. Our categorization of low and high task-correlation of datasets in this study was slightly arbitrary and limited by the number of scans analyzed. Increased reporting of task-correlation of motion, in addition to FD, would allow for further refinement in the usage of these combined metrics as a quality threshold or to guide denoising strategies.

Though our Conservative model demonstrated promising findings across scans, orthogonalization is a modeling strategy that should be applied thoughtfully, as it can confuse interpretation of activation results (Mumford et al., 2015). For example, orthogonalization of motion parameters to the task potentially attributes signal due to motion artifacts to task-related activation. This step can artificially inflate activation results, confounding proper interpretation of task-related brain activity. Therefore, in our Conservative model, instead of orthogonalizing the ME-ICA regressors to only the task, we orthogonalized the regressors to the entire core model, including motion parameters, end-tidal CO_2_, and Legendre polynomials. We aimed for the addition of ME-ICA regressors to aid in denoising, while not removing desired motor signal from being attributed to the core model. By also orthogonalizing these rejected nuisance regressors to all “accepted” (BOLD-weighted) ME-ICA components, we acknowledge that the true BOLD-weighted task activation will be present in one or more of those components, and that these data-driven representations of the signal of interest may be more accurate than our idealized regressor in the core model.

Here, we tested three methods of implementing ME-ICA nuisance regressors in subject-level models; however, other strategies and combinations with other denoising techniques may also be effective. For example, Beckers and colleagues (Beckers et al., 2023) found that combining ME-ICA denoising, using an approach like our Aggressive method, with an anatomical component-based correction method was highly effective for removing both focal and spatially diffuse noise in multi-echo fMRI data acquired at 7T. ME-ICA may also benefit from combination with physiological denoising, such as RETROICOR (Glover et al., 2000) and incorporation of heart rate variability (Chang et al., 2009) and respiratory volume per time (Birn et al., 2006), which could provide additional support in ME-ICA component classification. Integration with MARSS, another strategy that targets artifact resulting from head motion during multiband protocols (Tubiolo et al., 2024), may also be beneficial.

Lastly, while we used tedana version 24.0.2 in this analysis, tedana continues to undergo updates to improve reliability of available tools and implement new features. The task-correlated ME-ICA components identified and accepted in the Moderate model (Supplemental Figure 1) were highly associated with our signal of interest, indicating that ME-ICA component classification requires careful adjustment and quality control for the most accurate implementation by each study., New classification strategies may also improve this process; for example, Feng and colleagues found that tensor-ICA aided in classifying components by providing additional information about echo-time patterns (Feng et al., 2025). Studies that utilize tedana for ME-ICA component classification may benefit from a review of the latest updates and classification algorithms.

## Conclusion

In this study, we compared three methods of incorporating ME-ICA nuisance regressors in subject-level models of motor-task fMRI data. The Aggressive method involved simple inclusion of rejected ME-ICA components as nuisance regressors, the Moderate method involved exclusion of task-correlated ME-ICA components from the subject-level model, and the Conservative method involved orthogonalization of rejected ME-ICA components to the core subject-level model and accepted ME-ICA components. We tested these three methods in motor-task datasets from healthy and multiple sclerosis populations and during performance of hand-grasp, shoulder-abduction, and ankle-flexion motor tasks. We found that the Conservative model led to greater t-statistics, motor activation, and test-retest similarity in motor regions compared to the Aggressive model, particularly when amount and task-correlation of head motion was high and when the motor-task inherently produced lower signal (shoulder-abduction and ankle-flexion). We suggest that motor-task fMRI studies can implement the Conservative method to preserve signal of interest in subject-level activation maps.

## Author Contributions

Neha A. Reddy: Conceptualization, Methodology, Software, Formal analysis, Investigation, Data curation, Writing - original draft, Writing - review & editing, Visualization, Project administration. Michelle C. Medina: Conceptualization, Methodology, Software, Formal analysis, Investigation, Writing - review & editing. Jordan N. Northrop: Conceptualization, Methodology, Software, Formal analysis, Investigation, Writing - review & editing. Chendong Zou: Conceptualization, Methodology, Software, Formal analysis, Investigation, Writing - review & editing. Rebecca G. Clements: Conceptualization, Methodology, Writing - review & editing. Aravind Nehrujee: Writing - review & editing. Milap Sandhu: Methodology, Supervision, Writing - review & editing, Funding acquisition. Molly G. Bright: Conceptualization, Methodology, Writing - review & editing, Supervision, Project administration, Funding acquisition.

## Acknowledgements

This work was supported by the National Institute of Child Health and Human Development at the National Institutes of Health (R21HD108587, R03HD113915), the National Institute of Biomedical Imaging and Bioengineering at the National Institutes of Health (T32EB025766 to N.A.R.), the National Science Foundation (DGE-2234667 to R.G.C.), and the American Heart Association (25PRE1356822 to M.C.M.). The content is solely the responsibility of the authors and does not necessarily represent the official views of the National Institutes of Health. This work was supported by the Center for Translational Imaging at Northwestern University and through the computational resources and staff contributions provided for the Quest high performance computing facility at Northwestern University, which is jointly supported by the Office of the Provost, the Office for Research, and Northwestern University Information Technology.

