## Supplemental Material for "Impact of multi-echo ICA modeling decisions on motor-task fMRI analysis"

### Supplemental Materials

**Supplemental Table 1.** Scan parameters for functional scans acquired in each dataset.

| Dataset | Volumes | TR (s) | TE (ms) | FA (°) | MB factor | GRAPPA | Voxel size ($mm^3$) |
|---|---|---|---|---|---|---|---|
| **Healthy Hand** | 210 | 2 | 10.8/28.03/45.26/62.49/79.72 | 70 | 4 | 2 | 2.5x2.5x2 |
| **Healthy Shoulder** | 250 | 2.2 | 13.4/39.5/65.6 | 90 | 2 | 2 | 1.731x1.731x4 |
| **MSHand** | 258 | 1.5 | 10.6/27.83/45.06/62.29/79.52 | 70 | 4 | 2 | 2.5x2.5x2.5 |
| **MSFoot** | 324 | 1.5 | 10.6/27.83/45.06/62.29/79.52 | 70 | 4 | 2 | 2.5x2.5x2.5 |

A

| Dataset | # scans with task-correlated components / # total scans | % variance explained by task-correlated components |
|---|---|---|
| HealthyHand | 7 / 16 | 2.8 ± 1.1 |
| MSHand | 2 / 14 | 1.1 ± 0.1 |
| MSFoot | 4 / 14 | 1.3 ± 0.3 |
| HealthyShoulder | 8 / 28 | 5.6 ± 6.0 |

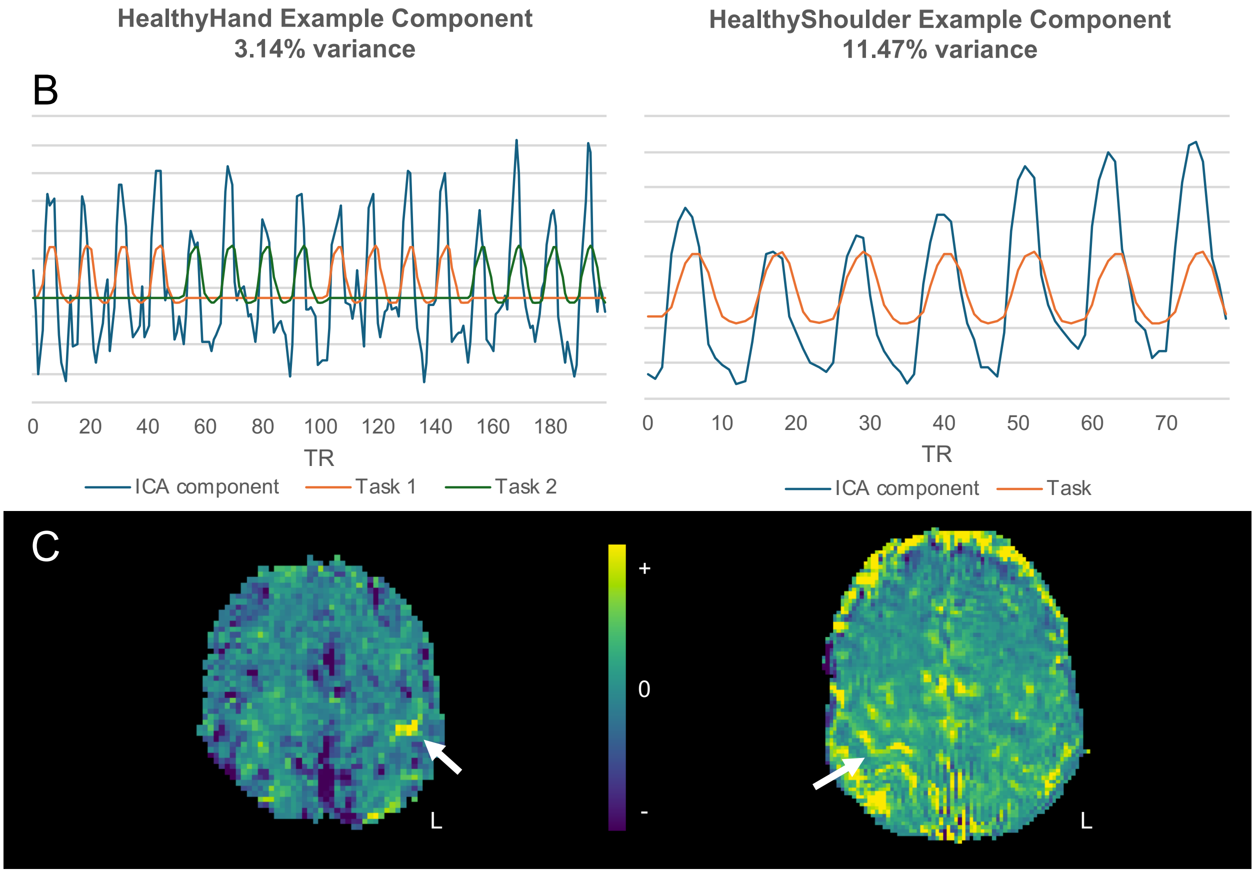


**Supplemental Figure 1.** Task-correlated components identified during ME-ICA classification were ‘accepted’ in the Moderate model. (A) Summary of scans with identified task-correlated components across datasets; in each of these scans, 1 or 2 components were identified. Two examples of task-correlated components are displayed, including (B) time course compared to task regressor(s) and (C) spatial maps. White arrows indicate regions of expected motor activity.

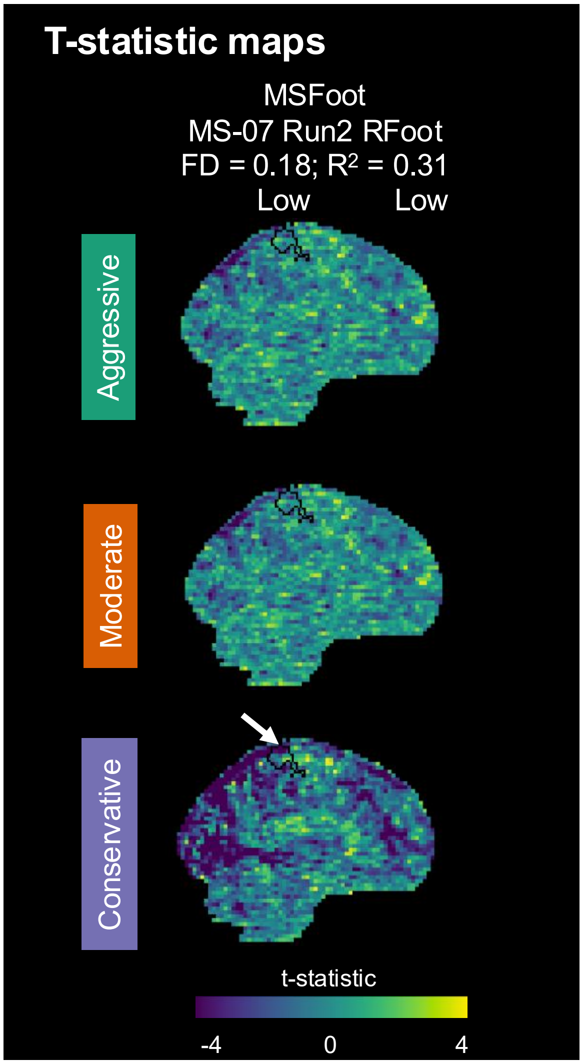


**Supplemental Figure 2.** Example MSFoot scan demonstrates artifactual negative t-statistics that are retained with the Conservative model. The primary motor cortex ROI is shown in black outline.

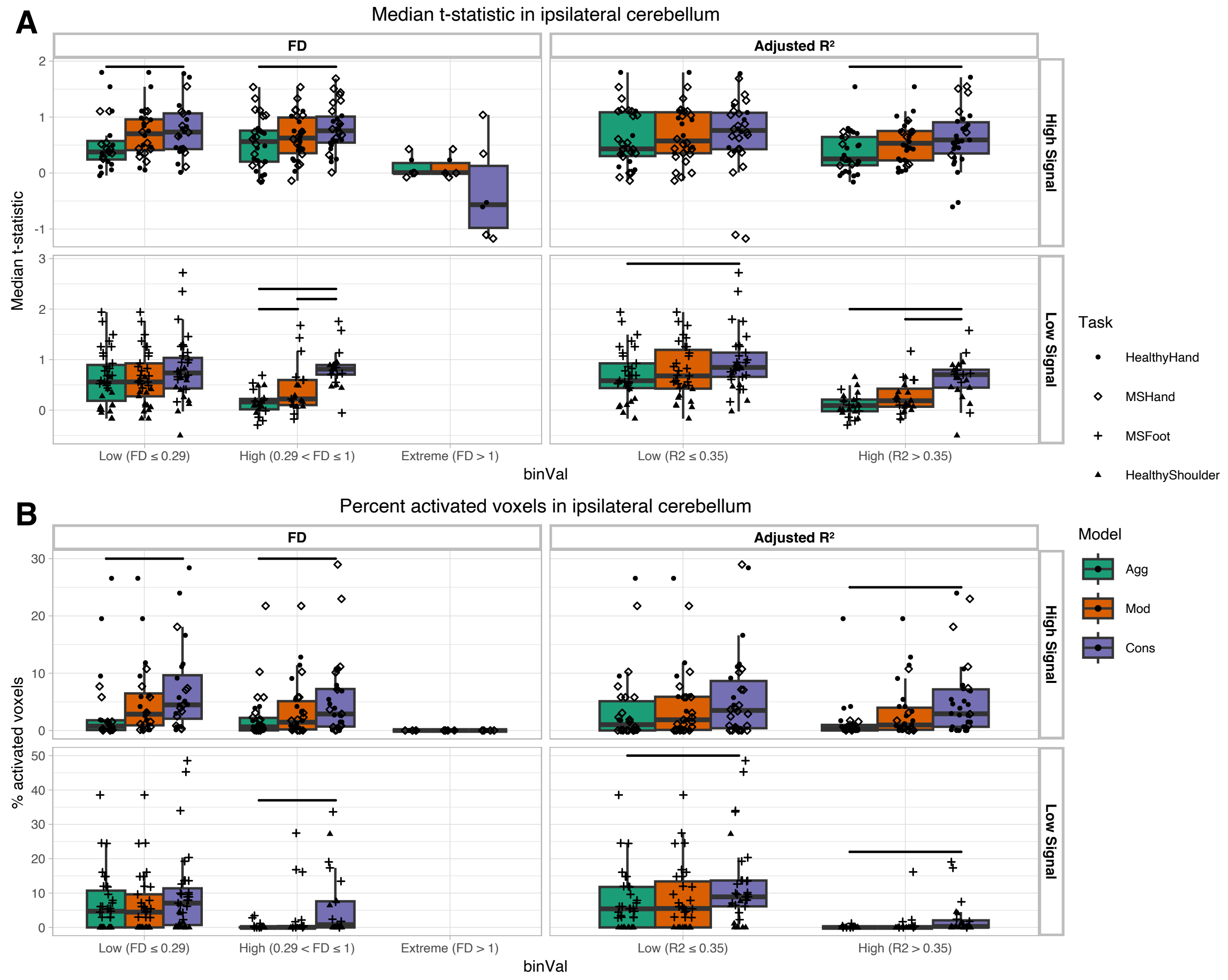


**Supplemental Figure 3.** Activation metrics calculated in significant group-level regions within the ipsilateral cerebellum. Regions were determined as the union of significant regions across all three models. (A) Median t-statistic and (B) Percent activated voxels were calculated for each scan. Horizontal bars indicate significant differences between models ($p < 0.05$, Bonferroni-corrected).

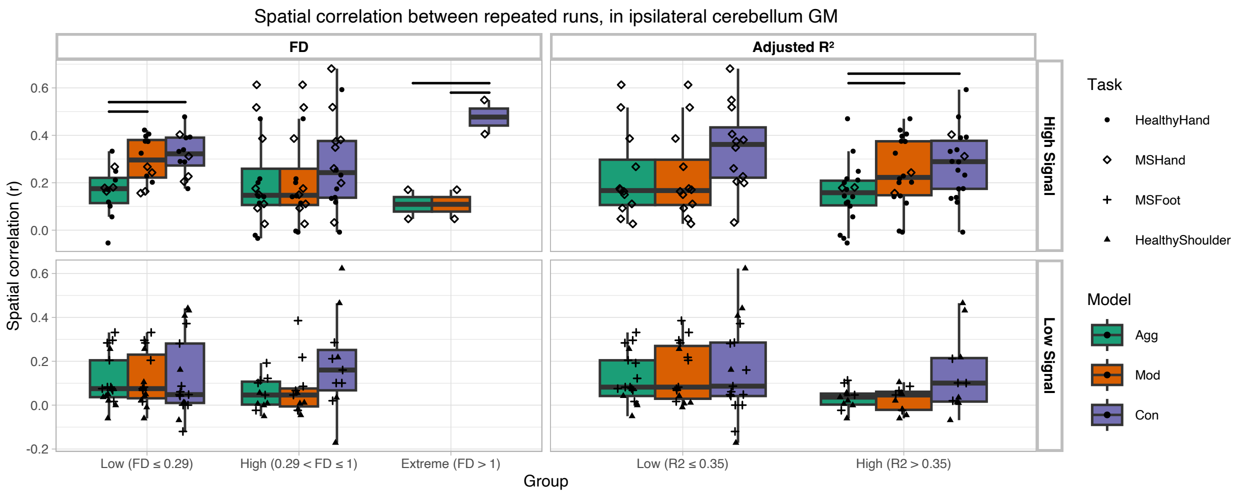


**Supplemental Figure 4.** Spatial correlation between runs 1 and 2 for each dataset and model, calculated in gray matter (GM) of the ipsilateral cerebellum. Horizontal bars indicate significant differences between models ($p < 0.05$, Bonferroni-corrected).
